# *Bacopa monnieri* phytochemicals regulate fibroblast cell migration via modulation of focal adhesions

**DOI:** 10.1101/2023.01.18.524521

**Authors:** Ravindra K. Zirmire, Dyuti Saha, Habibu Tanimu, Rania Zaarour, Deborah Bird, Prakash Cherian, Namita Misra, Aryasekhar Sanyal, Nita Roy, Colin Jamora

## Abstract

The *Bacopa monnieri* plant contains a large repertoire of active phytochemicals that have been used extensively in traditional medicine for the treatment of various complex diseases. More recently it has been shown to increase the wound healing rate in rats, though its mechanism of action is largely unknown. Here we investigated the cellular pathways activated by a methanol extract of *Bacopa monnieri* in human dermal fibroblasts, which play many critical roles in the wound healing program. Gene expression analysis revealed that *Bacopa monnieri* extract can enhance tissue repair by modulating multiple processes involved in the wound healing program such as migration, proliferation, and angiogenesis. We discovered that *Bacopa monnieri* extract can increase migration of fibroblasts via modulating the size and number of focal adhesions. *Bacopa monnieri*-mediated changes in focal adhesions are dependent on α5β1 integrin activation and subsequent phosphorylation of Focal Adhesion Kinase (FAK). Altogether our results suggest that *Bacopa monnieri* extract could enhance the wound healing rate via modulating fibroblast migration in the wound bed.

## INTRODUCTION

Wound healing comprises three distinct but overlapping phases. The initial phase is the inflammatory phase marked by the activation of resident immune cells and infiltration of circulating immune cells. The second phase is the proliferative phase wherein the proliferation and migration of fibroblasts, keratinocytes, and stem cells occur, in addition to the secretion of extracellular matrix (ECM) and angiogenesis. Finally, the wound-healing program culminates in the remodeling phase which comprises the removal of excess extracellular matrix and restructuring of cell-cell and cell-matrix interactions. Impairment of any of these phases can lead to delays in the kinetics of the wound-healing program. The phases of the wound healing program are the result of extensive and carefully orchestrated intercellular crosstalk. A major regulatory node of this intercellular network is the dermal fibroblasts that have been shown to impact all the phases of the wound healing program such as inflammation(1), angiogenesis(2), stem cell proliferation, and migration(3),(4) and protease release for tissue remodeling(5).

However, in various disease contexts, prolonged inflammation can lead to fibroblast senescence(6) and reduced wound healing activities. As a consequence of crippling fibroblast activity, the wound-healing program is significantly impaired. Thus, one method of restoring wound healing in chronic inflammatory conditions such as diabetes would be to repair fibroblast dysfunction. Herbal remedies have been proposed as a method to stimulate proliferation and differentiation of human mesenchymal stromal cells for cell therapies (7) and we investigated whether this same approach can be utilized for the promotion of fibroblast wound-healing activities. Extracts of *Bacopa monnieri* have been used in traditional medicine for the treatment of multiple diseases including Alzheimer’s disease(8), epilepsy(9), and anxiety and depression(10). In addition, it has been used for memory enhancement(11), as a nootropic agent(12), and interestingly for the enhancement of wound healing(13).

It has been shown that *Bacopa monnieri* extract, when administered orally, increases the wound closure rate in rats through its antioxidant activity(13),(14). It increased the levels of antioxidants such as GSH, SOD, and CAT whereas it reduced the levels of free radicals LPO and NO(13). Oxidants and free radicals are regulated by the inflammatory environment during the wound healing process(15). There was also a significant reduction in the inflammatory cells as well as inflammatory markers, increased neovascularization(13), and increased reepithelialisation(14). Altogether these results suggest that *Bacopa monnieri* extract affects inflammatory cells, endothelial cells, and keratinocytes to accelerate the wound healing process though the mechanisms and signaling pathways mediating these effects remain unknown. Given the importance of dermal fibroblasts in the cutaneous wound healing response, we investigated whether *Bacopa monnieri* extract can modulate the wound-healing activities of human fibroblasts and the mechanism of its action.

## MATERIAL AND METHOD

### Test substance

*Bacopa monnieri* extract was procured from Natural Remedies Pvt. Ltd, Bangalore under the brand name BacoMind™. It has the following bioactive constituents viz., bacoside A3 (>4.0% w/w), bacopaside I (>5.0% w/w), bacopaside II (>4.5% w/w), jujubogenin isomer of bacopasaponin C (>5.0% w/w), bacopasaponin C (>3.5% w/w), bacosine (>1.0% w/w), luteolin (>0.2% w/w), apigenin (>0.1% w/w) quantitated by HPLC.

### Cells

Human adult dermal fibroblasts (Lonza CC-2511) were cultured in DMEM media with (10% FBS, Pen-Strep, NEAA, and sodium pyruvate) in 5% CO2. All the experiments were performed with 3 biological replicates.

### RNA extraction, cDNA preparation, and real-time PCR

Human adult dermal fibroblasts were lysed in 1 ml Trizol reagent (Invitrogen, 10296028), and RNA was isolated according to the manufacturer’s protocol. cDNA was synthesized by PrimeScript cDNA synthesis kit (Takara, Cat No: 2680A) according to the manufacturer’s protocol. Quantitative PCR (qPCR) was carried out using Power SYBR Mix (Applied Biosystems, Thermo Fisher Scientific, Cat No: A25742) in a Bio-Rad CFX384 machine. GAPDH or TBP expression was used for normalization. Primers used for qPCR are listed below. PTGS2 forward primer) TCCAATGACTCCCAGTCTGAGGA, PTGS2 reverse primer TCAAAGGTCAGCCTGTTTAC, KIT forward primer TGTGTTGTCACCCAAGAGATT, KIT reverse primer CAATGAAGTGCCCCTGAAG, ARHGAP22 forward primer TACAGGGGCTGGTCACTGAG, ARHGAP22 reverse primer GTTCCGCAGCTTTATTTCCA, NEDD9 forward primer GCCTCTAGAAGCAAGTCCGC, NEDD9 reverse primer GGGAGGTGACAGCTAGTCCT.

### Statistics

A comparison of two groups to analyze effects on human adult dermal fibroblasts was made using the paired Student’s t-test. GraphPad Prism 5.02 was used for all the statistical analysis for all the experiments. Graphs represent mean ±SEM. P values less than 0.05 were considered to be significant.

### RNA Seq and data analysis

Adult human dermal fibroblasts were treated with vehicle control (water) or *Bacopa monnieri* for 12 hours and RNA was isolated as described previously. RNA sequencing was performed by MedGenome (Bangalore, India) and ∼45–55 million 150 bp paired-end reads were derived from three replicates of each treatment. The analysis was performed by using standard RNA Seq analysis pipeline. FASTQC tool (https://www.bioinformatics.babraham.ac.uk/projects/fastqc/) was employed for the QC of the raw sequencing reads. Low-quality reads at both ends were trimmed using “FASTX-TRIMMER” by using the parameters as “-f 11 -l 130” (http://hannonlab.cshl.edu/fastx_toolkit/index.html). Only the good-quality trimmed reads (120bp*2) were aligned with the human reference genome (hg38, UCSC version) using “HISAT2.”(16) The “Count” matrix was created using the “BAM” outputs using “HTSeq-count. (17) The differentially expressed genes were identified from the count matrix by using “DESeq2” R package.(17) Moreover, the significantly differentially expressed genes were used for Gene Ontology (GO) enrichment analysis using “DAVID” (18).

### Transwell migration assay

Transwells with 8 μm pore size (Corning, Cat. #3422) was used for transwell migration assay as described previously (19). 10µg/ml of *Bacopa monnieri* or vehicle control was added to the bottom compartment. The number of migrated cells on the lower compartment were stained with crystal violet and counted in three different fields.

### Cell proliferation assay

Human adult dermal fibroblasts were seeded at a density of 20000 cells per well in 24 well plates. Cells were cultured in 10% serum for 24 h, followed by treatment with vehicle control or *Bacopa monnieri* extract 5μg/ml in serum-free media. Cells were trypsinized and counted by using a hemocytometer on day 1, 2, 3.

### Cell viability assay

1000 human adult dermal fibroblasts were seeded per well of a 96 well plates and were allowed to attached completely for 24 hr in complete DMEM media. Post 24 hr cells were treated with vehicle control or *Bacopa monnieri* extract at doses of 5, 10, 15, 20, 25, and 30 µg/ml for 72 hr. The culture media was removed and cells viability was assessed using a 10% WST reagent (Roche, Cat. #5015944001) by incubating the cells for 2 hours and measuring the O.D. at 450nm.

### Immunofluorescence

For staining, cells were fixed with 4% paraformaldehyde (PFA) (Fisher Scientific, Cat. #50-980-487) on coverslips. Anti-integrin β1 antibody (12G10) (Abcam Cat. #ab30394) was used at a dilution of 1:200. For phospho-FAK (Y397) (Thermo Fisher, Cat. #44624g) staining, cells on the coverslip were fixed with chilled methanol at -20^0^C for 5 minutes and incubated with antibody at a dilution of 1:100. Alexa Fluor 488 or Alexa Fluor 568–labeled secondary antibodies (Jackson Immuno Research) was used at a dilution of 1:200. Fluorescence intensity for anti-integrin β1 antibody (12G10) and pFAK was quantitated using Image J with the following formula: Corrected Total Cell Fluorescence (CTCF) = Integrated Density – (area of selected cell x mean fluorescence). The size, shape, and number of focal adhesions were calculated by Image J software (National Institutes of Health, Bethesda, USA) by using the “Manual” freehand tool.

### Scratch migration assay

Migration in a scratch assay was performed as described (20). Briefly, 50000 cells were seeded in each well in 24 well plates. The cells were grown to a confluent monolayer in complete DMEM media. The cells were then washed with PBS and a “wound” was introduced by scratching the plate with a sterile 1ml pipette tip. The debris was then removed by washing the cells with PBS. The cells were treated with vehicle control or 5µg/ml *Bacopa monnieri* extract. The scratch closure was analyzed by imaging the scratch daily until the scratch was closed. The open area of the scratch was analyzed by Image J freehand tool. The scratch closure was plotted as % open wound.

### Random migration assay

5000 fibroblasts were added to each well of a 24 well dish and allowed to adhere for 24 hours in complete media. Cells were then treated with vehicle control or 5µg/ml *Bacopa monnieri* extract in serum free media. Live cell imaging was performed with a phase-contrast microscope setting on IX 83 microscope with a motorized stage. 5 different fields were imaged at 15 minutes time intervals for 12 hours. Single cells were tracked by tracking the nucleus of the individual cell by ImageJ software using the “Manual Tracking” plugin (Fabrice Cordelières, Institut Curie, Orsay, France). Each experiment was repeated three times independently. Further analysis for distance, velocity, and directionality was performed by using the Chemotaxis and Migration Tool V2.0 from Ibidi. At least 50 cells were analyzed per condition.

## RESULTS

### *Bacopa monnieri* extract increases fibroblast migration

To test the effect of *Bacopa monnieri* extract on primary human dermal fibroblasts, we first determined the maximal tolerable concentration by measuring cell viability in the presence of the extract (Supplementary Figure 1A). The maximum tolerable concentration was found to be 10μg/ml. To take an unbiased approach to decipher the effect of *Bacopa monnieri* extract, RNA sequencing analysis was performed on adult fibroblasts treated with *Bacopa monnieri* extract for 12 hours. Interestingly, gene cluster analysis revealed that *Bacopa monnieri* extract modulated several genes associated with multiple processes of the wound-healing program (Figure 1A). Though the positive regulation of cell proliferation was the top biological process, *Bacopa monnieri* extract-treated fibroblasts did not display an increase (Supplementary Figure 1B) or decrease (Supplementary Figure 1C) in the proliferation rate compared to control-treated cells.

**Figure 1.**
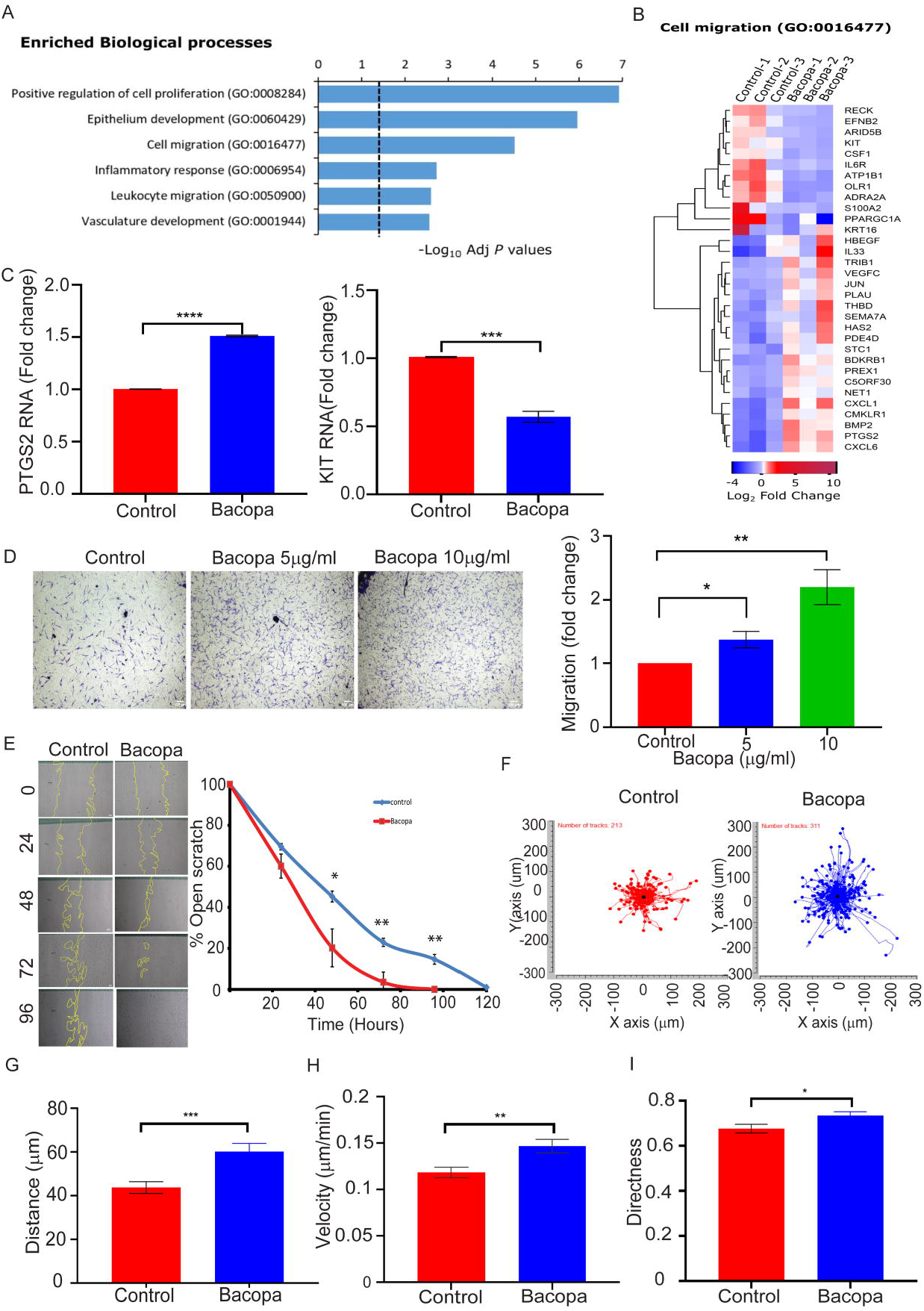
*Bacopa monnieri* extract increases human dermal fibroblast migration. A) Enriched biological processes after RNA sequencing analysis of fibroblasts treated with *Bacopa monnieri* extract. B) Heat map of genes associated with cell migration in cells treated with *Bacopa monnieri* or vehicle control (water). C) qPCR analysis of PTGS2 and KIT gene expression. D) Transwell migration of cells in the presence of *Bacopa monnieri* or vehicle control was analysed by crystal violet staining (left panel) and quantified (right panel). E) Scratch migration of cells in the presence of Bacopa or vehicle control (left panel) and quantification (right panel). p values were calculated using paired Student’s t test. *p<=0.05, **p<0.01, ***p<0.001, ****p<0.0001, (n=3 biological replicates). F) Quantification of 2D random migration of fibroblasts treated with *Bacopa monnieri* extract or vehicle control. Analysis of Distance (G), Velocity (H) and Directness (I) of cell migration of individual fibroblasts in the presence of Bacopa *monnieri* extract (n=311 cells) or vehicle control (n=213 cells). p values were calculated using unpaired Student’s t test. *p<=0.05, **p<0.01, ***p<0.001

On the other hand, a major biological process that was deemed to be affected by *Bacopa monnieri* exposure is cell migration (Figure 1A and B). For instance, quantitative PCR confirmed that the pro-migratory gene PTGS2 (21) is up-regulated, while the migration inhibiting gene ADRA2A (22) is down-regulated (Figure 1C). To determine whether the changes in gene expression are manifested in the biological activity of the dermal fibroblasts, we performed a transwell migration assay. Primary human dermal fibroblasts were seeded in the top compartment of the transwell membrane and *Bacopa monnieri* extract or vehicle control was added to the media of the lower compartment. 12 hours post-incubation, cells were fixed and the cells which had migrated from the top of the transwell to the bottom were counted. Interestingly, it was observed that *Bacopa monnieri* extract increased the chemotactic migration of fibroblast in a concentration-dependent manner (Figure 1D). In order to test if *Bacopa monnieri* extract can increase migration in a two-dimensional migration platform, a scratch wound assay was performed. Cells were cultured to form a confluent sheet and treated with mitomycin C to block proliferation. A “wound” was induced by using a pipet tip to scratch the cells and form a gap in the cell sheet. The scratch wound was monitored for the rate of closure by imaging and quantifying the wound area every 24 hours. It was observed that *Bacopa monnieri* extract substantially increased the wound closure rate of cells (Figure 1E). Since *Bacopa monnieri* extract increased the directional migration of cells in both a 3D and 2D assay, we investigated whether it also promoted the general motility of the fibroblast. Analysis of random migration was performed by the live tracking of the nuclear location of individual, underconfluent cells with brightfield microscopy in the presence of *Bacopa monnieri* extract or vehicle control for 12 hours (Figure 1F). The motility analysis of single cells was performed with the ImageJ manual tracker and revealed that *Bacopa monnieri* extract significantly increased the average distance moved by the cell from its starting point at the beginning of the analysis (Figure 1G). In addition, the velocity of the *Bacopa monnieri* treated cells was elevated relative to fibroblasts treated with vehicle control (Figure 1H). Interestingly, it was observed that *Bacopa monnieri* extract also increased the directionality of the migration (Figure 1I). Overall, this data indicates that *Bacopa monnieri* extract increased the basic motility of the fibroblasts as well as their directional migration.

### *Bacopa monnieri* extract treatment modulates focal adhesions in human dermal fibroblasts

To elucidate the mechanism by which *Bacopa monnieri* extract increases human dermal fibroblast migration, the transcriptome profile was analyzed for enriched cellular components. Interestingly, it was observed that the *Bacopa monnieri* extract modulated several genes associated with cell substratum adhesion and genes associated with structural components of focal adhesions (Figure 2A). We confirmed the changes in focal adhesion associated genes by RT-PCR and it validated the upregulation of ARHGAP22, a member of the RhoGAP family of proteins with pro-migratory effects (23) and downregulation of NEDD9, which stabilizes focal contacts and suppresses migration (24) (Figure 2B). The focus on focal adhesions is particularly relevant since their size and shape have been shown to uniquely predict the rate of cell migration (25), (26) To assess if *Bacopa monnieri* extract also affects the size of focal adhesions, vinculin (a structural component of focal adhesions) was characterized as a proxy for focal adhesion morphology. The size and shape of focal adhesions were quantified using Image J manual measuring with the “freehand selection” tool. Interestingly, it was observed that the *Bacopa monnieri* extract-treated fibroblasts exhibited reduced focal adhesion size throughout the cell (Figure 2C), and an altered shape from elongated to a more spherical morphology (Figure 2D). Additionally, it was found that the decreased size and increased roundness of the focal adhesions in *Bacopa monnieri* extract-treated cells were compensated by an increased number of focal adhesions (Figure 2E).

**Figure 2.**
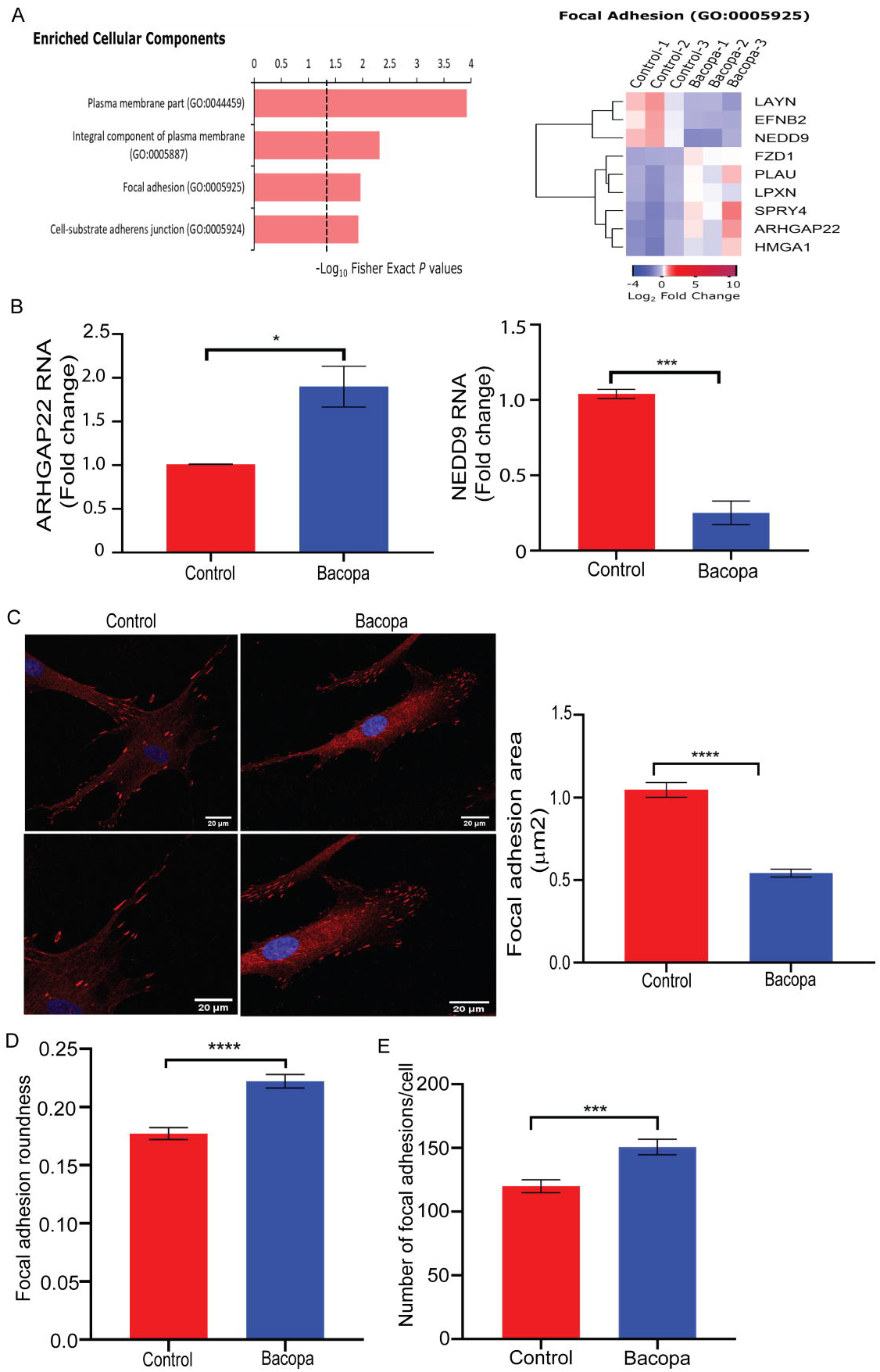
*Bacopa monnieri* extract alters focal adhesions in fibroblasts. A) Cellular component analysis from RNA sequencing of fibroblasts treated with *Bacopa monnieri* extract (left panel). Heat map of genes associated with focal adhesions (right panel). B) qPCR analysis of ARHGAP22 and NEDD9 gene expression. C) Vinculin staining (red) in human dermal fibroblast treated with *Bacopa monnieri* extract or vehicle control, DAPI (blue) marks nuclei (left panel) and quantification of focal adhesion area (right panel) scale bar 20μm. Analysis of focal adhesions roundness (D) and number per cell (E). p-values were calculated using unpaired Student’s t test. *p<=0.05, **p<0.01, ***p<0.001, vehicle control n=138 cells, *Bacopa monnieri* extract n=144 cells.

### *Bacopa monnieri* extract activates α5β1 integrin

Given that integrins regulate the de novo assembly of focal adhesions (27) and have a major impact on the migratory behaviour of cells (28), (29), we assessed whether *Bacopa monnieri* extract regulates these processes at the level of integrin receptors. We focused on integrin α5β1, which is found on human dermal fibroblasts and is known to modulate their migration (29). The status of α5β1 integrin activation was assessed by immunostaining the *Bacopa monnieri* extract-treated cells with an active integrin antibody (12G10), which detects the integrin only when it is in its active conformation. Corrected total cell fluorescence (CTCF) analysis by Image J revealed that *Bacopa monnieri* extract increased the activation of integrin by ∼50% compared to the vehicle control (Figure 3A). To investigate whether the *Bacopa monnieri* -mediated activation of α5β1 is responsible for the enhanced cellular migration, we inhibited the integrin with the RGD peptide that competitively binds to their ECM binding site (30). Pretreatment of cells with a high concentration (500uM) of RGD peptide was done followed by a trans-well migration assay in the presence of *Bacopa monnieri* extract in the lower chamber. Interestingly, it was observed that the addition of RGD peptide effectively blocked the *Bacopa monnieri* extract-induced chemotaxis (Figure 3B). To test if inhibition of integrin signaling by RGD could inhibit the decrease in the size of focal adhesions caused by *Bacopa monnieri*, human dermal fibroblasts were treated with extract in the presence of 500uM of the RGD peptide. Cells were immunostained for vinculin and analyzed for the effect on focal adhesion size. It was observed that RGD peptide increased the size of focal adhesions significantly compared to the vehicle control. Interestingly, inhibition of integrin activation in *Bacopa monnieri* extract-treated cells blocked the reduction in the size of focal adhesions (Figure 3C). Inhibition of integrin activity could also reduce the roundness of focal adhesions in *Bacopa monnieri* extract-treated fibroblasts (Figure 3D). Additionally, the RGD peptide significantly inhibits the increase in the number of focal adhesions per cell compared to *Bacopa monnieri* extract alone (Figure 3E). Taken together it was observed that *Bacopa monnieri* extract increases fibroblast cell migration by activating integrin signaling.

**Figure 3.**
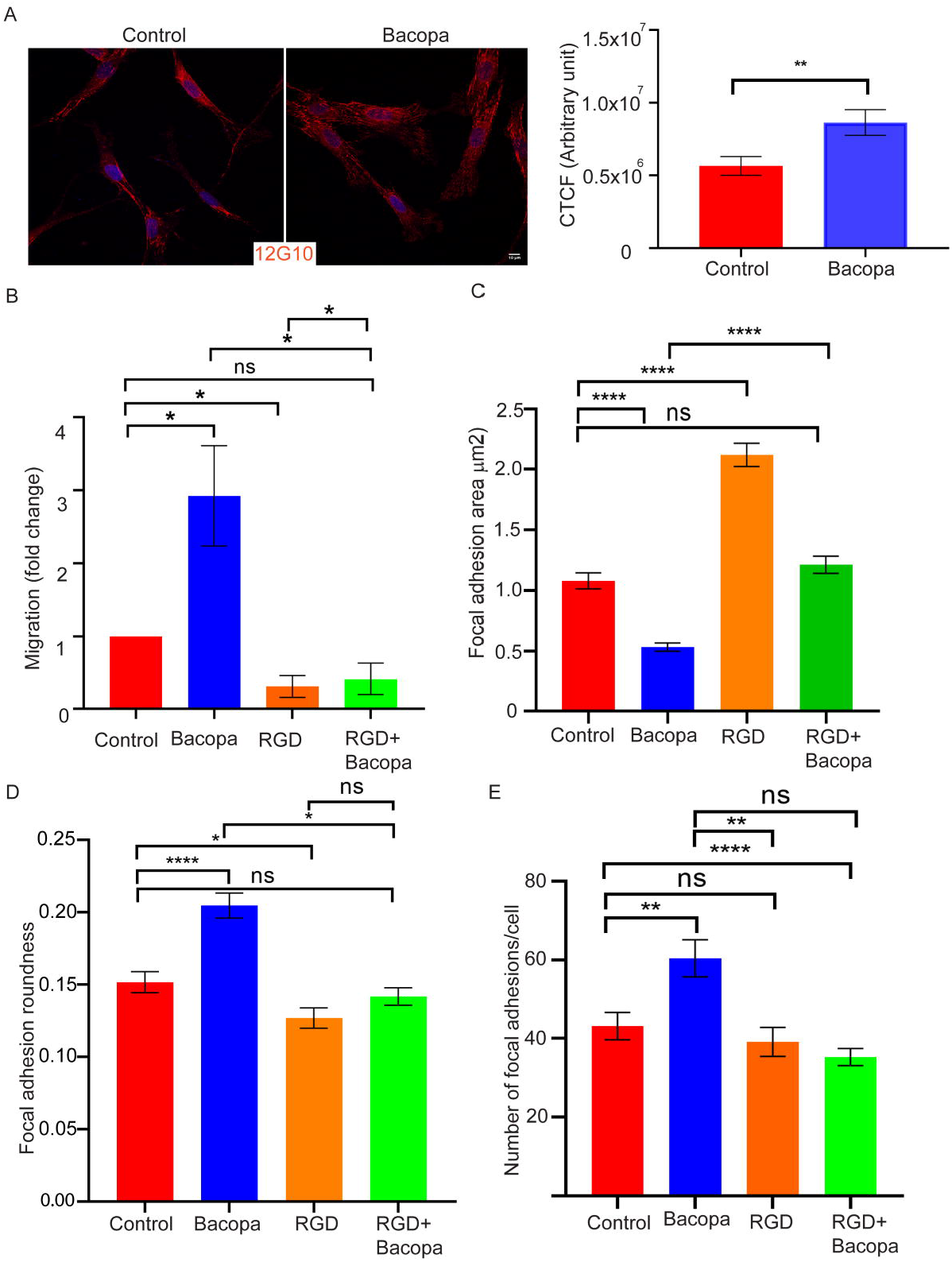
α5β1 integrin activation mediates *Bacopa monnieri* extract-induced migration. A) Immunofluorescence staining for active integrin 12G10 (red) in cells treated with *Bacopa monnieri* or vehicle control (left panel) and corrected total cellular fluorescence (CTCF; right panel). Scale bar 10μm B) Quantification of transwell migration of cells in the presence of vehicle control, *Bacopa monnieri* extract, RGD peptide, or *Bacopa monnieri* extract + RGD peptide. p-values were calculated using unpaired Student’s t test. *p<=0.05, (n=3 Biological replicate). C) Quantification of focal adhesion area or roundness (D) and number (E) in control (n=67 cells), *Bacopa monnieri* treated (n=67 cells), RGD peptide treated (n=41 cells), or *Bacopa monnieri* extract + RGD peptide treated cells (n=41 cells) in three independent experiments. p values were calculated using unpaired Student’s t test. *p<=0.05, **p<0.01, ***p<0.001, ****p<0.0001.

### *Bacopa monnieri* extract activates focal adhesion kinase

One mechanism by which integrin signaling mediates migration is through focal adhesion kinase (FAK) (31). The status of FAK activation was assessed by immunostaining the *Bacopa monnieri* -treated cells with pFAK antibody which selectively detects the phosphorylated (active) form of FAK. Interestingly it was observed that *Bacopa monnieri* extract increased the phosphorylation of FAK by ∼30% (Figure 4A). To investigate if FAK activity mediates *Bacopa monnieri* extract-induced cellular migration, we inhibited the FAK phosphorylation with 5µM of the inhibitor PF-573228. Interestingly, it was observed that the addition of FAK inhibitor blocked the *Bacopa monnieri* extract induced transwell migration to control levels (Figure 4B). Likewise, we investigated whether inhibition of FAK signaling can abrogate the reduction in the size of focal adhesions caused by *Bacopa monnieri* extract. Human dermal fibroblasts were treated with extract in the presence of FAK inhibitor and focal adhesions were visualized with an antibody against vinculin. Interestingly, while FAK inhibitor alone did not affect focal adhesion size relative to control, the inhibitor restored the focal adhesions to normal size in *Bacopa monnieri* extract-treated cells (Figure 4C). Similarly, inhibition of FAK phosphorylation inhibits the increased roundness of focal adhesions in *Bacopa monnieri* extract-treated fibroblasts (Figure 4D). In addition, the restoration of focal adhesion morphology by FAK inhibitor also resulted in the normalization of focal adhesion number in *Bacopa monnieri* extract-treated cells (Figure 4E).

**Figure 4.**
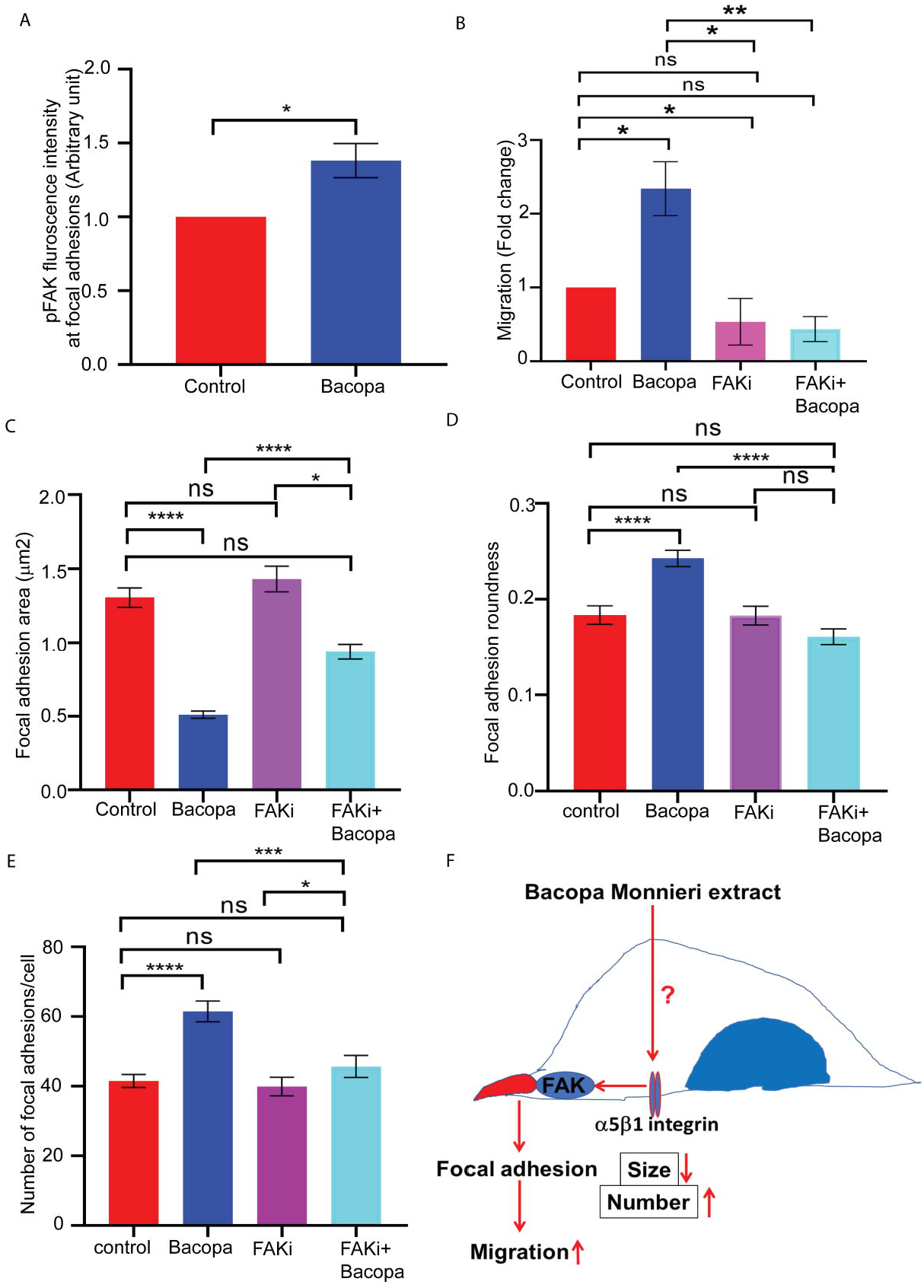
Focal adhesion kinase activation mediates *Bacopa monnieri* extract induced migration. A) Fluorescence intensity of phospho FAK in human dermal fibroblasts treated with *Bacopa monnieri extract* or vehicle control. B) Quantification of transwell migration of cells in the presence of vehicle control, *Bacopa monnieri* extract, FAK inhibitor, or *Bacopa monnieri* extract + FAK inhibitor. p-values were calculated using unpaired Student’s t test. *p<=0.05, (n=3 Biological replicate). (C) Quantification of focal adhesion area or roundness (D) and number (E) in control (n=41 cells), *Bacopa monnieri* treated (n=67 cells), FAK inhibitor treated (n=41 cells), or FAK inhibitor + *Bacopa monnieri* extract treated cells (n=61 cells) in three independent experiments. p values were calculated using unpaired Student’s t test. *p<=0.05, **p<0.01, ***p<0.001, ****p<0.0001.

## DISCUSSION

Altogether the data presented suggests a mechanism by which *Bacopa monnieri* extract can induce the migration of dermal fibroblast. Primary human dermal fibroblasts exposed to *Bacopa monnieri* extract can activate integrin α5β1 which subsequently leads to the activation of focal adhesion kinase (FAK). The active kinase leads to a decrease in the size and increase in the number of focal adhesions. This data is in line with reports that smaller focal adhesions correlate with a higher migratory capacity of the cell (25), (32). Consequently, this model partly explains the mechanism by which *Bacopa monnieri* extract can accelerate wound healing that was observed in vivo (13). By stimulating the migration of resident dermal fibroblasts, the *Bacopa monnieri* extract can drive the recruitment of these cells into the wound bed where they play important roles in regulating the tissue repair response such as inflammation, angiogenesis, and ECM production/remodeling. Deciphering the mechanism of action of *Bacopa monnieri* extract fills an important gap in the field of herbal remedies where the cellular processes modulated by phytochemicals are largely undetermined.

Though the use of herbal remedies generally lacks an in-depth mechanistic understanding, this method to promote physiological processes or correct pathological scenarios has various advantages over synthetic drugs. The latter has several drawbacks such as higher side effects and cost compared to herbal medicine (33). Moreover, herbal remedies are generally more widely available and non-toxic for therapeutic uses and are based on a more holistic approach rather than targeting specific symptoms of a disease.

The application of *Bacopa monnieri* extract in the wound healing context illustrates this holistic approach to restoring homeostasis. There are multiple reasons for the impairment of wound healing which may include local, systemic factors, and reduced tissue growth factors. Local factors comprise tissue maceration, foreign bodies, biofilm, hypoxia, ischemia, and wound infection. Systemic factors comprise diabetes, malnutrition, and other chronic organ diseases. In general, it is not possible to remove these factors completely even in good clinical practices (6). In addition to local and systemic factors, reduced levels of tissue-related growth factors increased proteolytic enzymes, and increased inflammatory mediators, also participate in a delayed wound healing program (34). As a result of the multi-tiered defects that lead to chronic, non-healing wounds many processes within the wound healing program are perturbed. We have now shown that the pro-migratory effect of *Bacopa monnieri* extract on human dermal fibroblasts complements the reported holistic effect of this herbal remedy that also increases the wound closure rate in rats, counteracts free radicals and reduces inflammation.

## ACKNOWLEDGEMENTS

The authors would like to thank Jamora lab members for critical review of the work and insightful discussions, and the inStem/NCBS Central Imaging and Flow Cytometry facility for generating the confocal images. This work was supported by grants from L’Oreal, the Department of Biotechnology, Government of India (BT/PR8738/AGR/36/770/2013), and core funds from Institute for Stem Cell Science and Regenerative Medicine (inStem) to CJ. RZ was supported by a DBT JRF fellowship, RFZ was supported by a postdoctoral fellowship from FIRC Institute for Molecular Oncology (IFOM), HT was supported by Dr. Bindu O.S (Jain University).

## LIST OF ABBREVIATIONS

FAK: (Focal adhesion kinase);
ECM: (Extra cellular matrix);
GSH: (Glutathione);
SOD: (Superoxide dismutases);
CAT: (catalase);
LPO: (Lipid peroxidation);
NO: (Nitric oxide).

## FIGURE LEGENDS

**Supplementary Figure 1. *Bacopa monnieri* extract does not affect cell proliferation rate.** A) Maximum tolerable dose was determined by WST assay. B) Proliferation assay with WST after treating the cells with *Bacopa monnieri* extract or vehicle control. C) Proliferation inhibition assay with WST after treating the cells with *Bacopa monnieri* extract or vehicle control. p values were calculated using paired Student’s t test. *p<=0.05, **p<0.01, ***p<0.001, three independent biological replicates.

**Supplementary Figure 2.** Bacopa monnieri extract activates focal adhesion kinase. A) Immunofluorescence staining for p-FAK in the cells treated with Bacopa monnieri extract of vehicle control. Scale bar 10μm.

