## Supplementary figures and images for "*Bacopa monnieri* phytochemicals regulate fibroblast cell migration via modulation of focal adhesions"

### Supplementary file 1

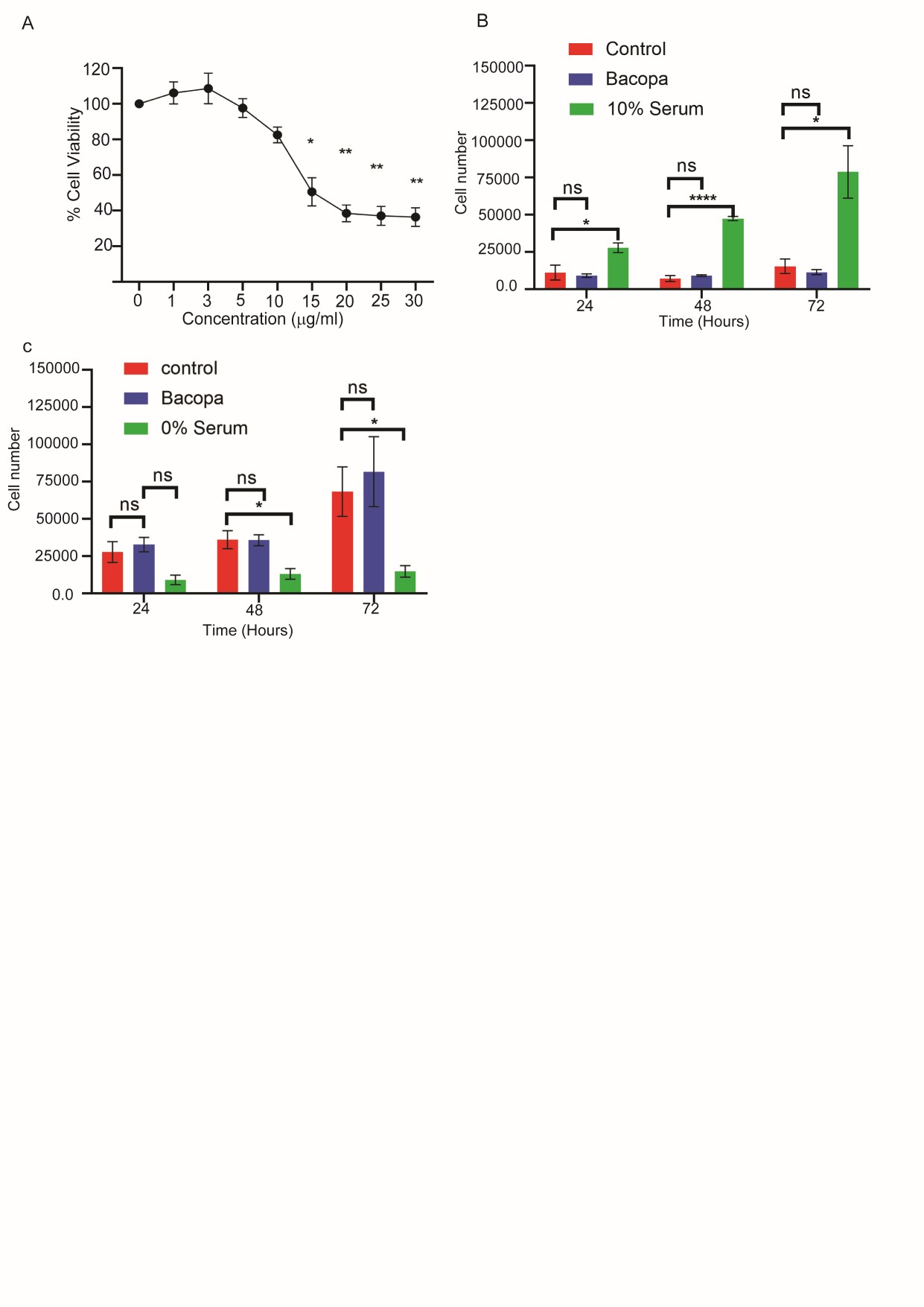

### Supplementary file 1

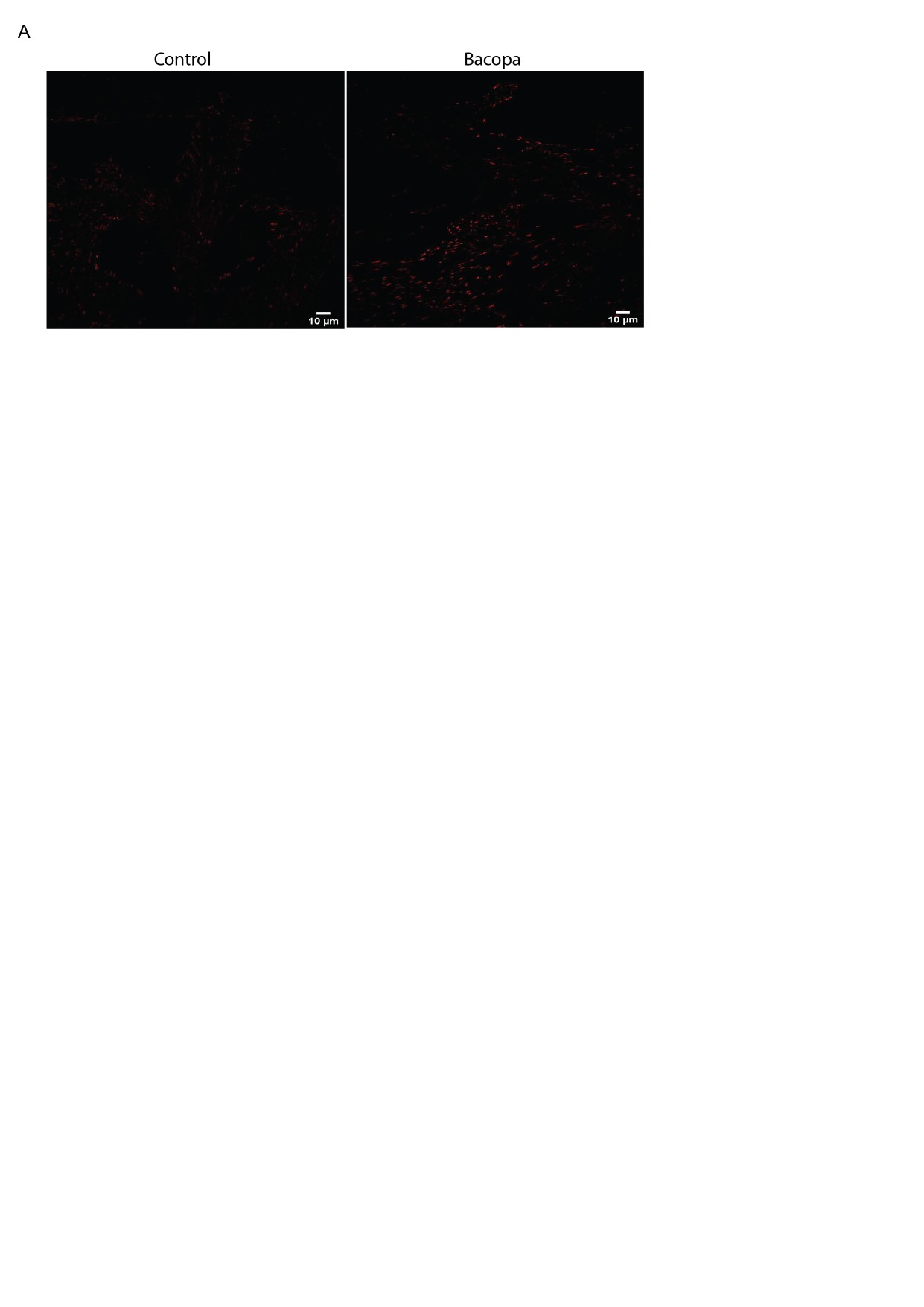
